# Transient ATM inhibition enhances knock-in efficiency in hematopoietic stem cells by attenuating the DNA damage response

**DOI:** 10.1101/2025.09.03.673991

**Authors:** Munkh-Erdene Natsagdorj, Hiromasa Hara, Kohei Iida, Yuji Kashiwakura, Tsukasa Ohmori, Yutaka Hanazono, Fumio Nakahara

## Abstract

Precise genome editing in hematopoietic stem cells (HSCs) offers great potential for treating inherited blood disorders, but low knock-in (KI) efficiency, due to HSC quiescence and a preference for non-homologous end joining (NHEJ) and DNA damage-induced apoptosis, remains a major barrier. Here, we demonstrate that transient inhibition of Ataxia-Telangiectasia Mutated (ATM) kinase markedly enhances KI efficiency in mouse HSCs genome-edited with Cas9/RNP and AAV donor DNA. Phosphoproteomic analysis and capillary western blotting revealed that ATM inhibition suppressed the Cas9-AAV-induced ATM activation and subsequent DNA damage response, reduced p53-dependent apoptosis and preserved knock-in competent cells. In transplantation experiments, ATM inhibition preserved long-term engrafting genome-edited HSCs, increasing their frequency from ∼0.3% to ∼40% in secondary recipients - a >100-fold enhancement compared to untreated cells. Furthermore, in an X-SCID mouse model, ATM inhibition enhanced KI efficiency and restored expression of IL-2 receptor γ chain (CD132). These strikingly novel findings highlight transient ATM inhibition as a powerful and clinically relevant approach to enhance KI-mediated genome editing in HSCs, while preserving their long-term repopulating capacity.

**Key points:**

- ATM inhibition enhances knock-in efficiency in mouse hematopoietic stem cells
- ATM inhibition suppresses Cas9-AAV-induced overactivation of ATM and subsequent p53-dependent apoptosis.

## Introduction

Hematopoietic stem cells (HSCs) are defined by their capacity for self-renewal and multilineage differentiation and are essential for the lifelong maintenance of the hematopoietic system^1,2^. Allogeneic HSC transplantation (HSCT) remains the standard treatment for a broad range of hematological malignancies and inherited blood disorders, including severe combined immunodeficiency (SCID), β-thalassemia, and sickle cell disease (SCD)^3–5^. Despite its curative potential, the clinical application of allogeneic HSCT has major limitations, notably the shortage of HLA-matched donors and the risk of graft-versus-host disease^6,7^. To overcome these challenges, gene therapy approaches have been developed to enable autologous correction of genetic defects^8–10^. However, most current gene therapy strategies rely on viral vector-mediated random integration of therapeutic transgenes, which is associated with variable expression and the potential for insertional mutagenesis^11^. In contrast, *ex vivo* genome editing followed by autologous transplantation offers a more precise alternative, allowing targeted gene modification of mutations^12^. The recent clinical success of Casgevy^®^ —a CRISPR/Cas9-based genome editing therapy that disrupts the erythroid-specific enhancer of *BCL11A* via non-homologous end joining (NHEJ) to induce fetal hemoglobin expression—has demonstrated the proof of concept and shown the therapeutic benefit of this approach in patients with β-thalassemia and SCD^13,14^. While Casgevy^®^ represents an innovative achievement, it relies on a gene disruption strategy specific to disorders with known repressor elements and compensatory pathways. For the broader range of monogenic hematologic diseases, precise gene correction at endogenous loci will be required, necessitating efficient knock-in (KI)-based genome editing^15–17^. KI is commonly achieved via Homology-directed repair (HDR), the pathway required for precise gene correction^18^. However, the efficiency of KI in HSCs remains challenging due to their quiescent state and DNA repair preferences for NHEJ over HDR, limiting the broader application of genome editing as a generalizable therapeutic platform. To overcome the limitation of low KI efficiency in HSCs, various strategies have been explored to enhance KI efficiency. These include *ex vivo* culture systems to stimulate HSC proliferation^19^, or pharmacological inhibition of NHEJ factors such as DNA Ligase IV^20,21^, DNA-dependent protein kinase catalytic subunit (DNA-PKcs)^22^ or the use of an inhibitory peptide for p53-binding protein 1 (53BP1)^23^, which compete with HDR and favor error-prone repair. While these approaches have shown some success in improving editing efficiency, concerns remain regarding their potential effects on HSC function, including stem cell homeostasis and long-term engraftment capacity. Therefore, alternative strategies that promote KI without impairing the stem cell properties of HSC are needed to advance genome editing for broader therapeutic application.

Ataxia-telangiectasia mutated (ATM) kinase functions as an early sensor of DNA double-strand break (DSB) and serves as a key regulator of the DNA damage response (DDR), coordinating the decision between DNA repair and apoptosis. Upon activation, ATM phosphorylates a broad range of substrates, including p53, H2AX, and 53BP1, which collectively influence cell fate and DNA repair pathway choice^24,25^. In our previous work using a mouse embryonic stem (ES) cell model, we found that the introduction of linear donor DNA—regardless of strand configuration—can lead to excessive ATM activation. This overactivation triggered apoptosis or biased repair toward NHEJ, thereby limiting KI efficiency. Importantly, transient inhibition of ATM reduced this apoptotic response and suppressed NHEJ activity, at least in part through downregulation of DNA-PKcs signaling. These effects resulted in a significant enhancement of KI efficiency in cell lines when using Cas9 as ribonucleoprotein (RNP) in combination with AAV donor templates (M.N., H.H., Hideki Uosaki, F.N., Makoto Inoue and Y.H., manuscript submitted June 2025). Expanding on these findings, we sought to translate this strategy to primary hematopoietic stem cells. Given the unique DNA repair feature and sensitivity of HSCs to genotoxic stress, we investigated whether transient ATM inhibition could similarly modulate the DDR to favor KI without compromising HSC stemness.

To evaluate whether transient inhibition of ATM could enhance KI efficiency in HSCs, we combined Cas9/RNP and single-stranded AAV donor DNA with ATM inhibition in *ex vivo* cultured mouse HSCs. We then investigated the impact of this approach on genome editing efficiency, DDR signaling, and functional stem cell activity using transplantation-based assays and phosphoproteomic profiling. This study was designed to assess whether modulating DDR pathway activation through the transient ATM inhibition could provide a generalizable strategy to improve the precision and therapeutic potential of HSC genome editing.

## Materials and Methods

### Mice

C57BL/6J (CD45.2) mice (Japan SLC) and C57BL/6J (CD45.1) mice (kindly provided by Dr. Atsushi Iwama) were used for genome editing and transplantation assays. Previously generated X-linked severe combined immunodeficient (X-SCID) mice were used for treatment of disease model^26^. All animal studies were conducted in accordance with the guidelines set forth by the Institutional Animal Care and Concern Committee at Jichi Medical University and were approved by the committee. All efforts were made to ensure the animal welfare and to minimize pain and distress throughout the experimental procedures.

### Preparation of mouse Lineage^-^ and c-Kit^+^ cells

To obtain bone marrow cells for use in this study, tibias and femurs were extracted from 8-week-old mice, and the cells were isolated. To remove erythrocytes, the cells were treated with 1×RBC lysis buffer (Pluriselect). Magnetic beads were then used to isolate Lineage^-^ and c-Kit^+^ (LK) cells using Direct Lineage Depletion cocktail and mouse CD117 microbeads (Miltenyi Biotech, Germany) on autoMACS separator.

### Cell culture

Mouse 32D cells were cultured in RPMI-1640 medium (Gibco) containing 10% Fetal Bovine Serum (Gibco), 1 µg/ml Mouse IL-3 (Peprotech), 1 × Glutamax (Gibco) and 1 × penicillin streptomycin (Gibco). The cells were cultured at 37°C in an atmosphere containing 5% CO2 and medium was changed every 2-3 days.

*Hes1*-expressing Common Myeloid Progenitor (Hes1-CMP) cells were cultured in IMDM medium (Gibco) containing 10% Fetal Bovine Serum (Gibco), 1 µg/ml Mouse IL-3 (Peprotech), 1 × Glutamax (Gibco) and 1 × P/S (Gibco). The cells were cultured at 37°C in an atmosphere containing 5% CO2 and medium was changed every 2-3 days.

*Ex vivo* separated Lineage^-^ and c-Kit^+^ (LK) cells from C57BL/6J mice were cultured in HemEx-Type 9A (Cell Science & Technology Institute, Inc) containing 100 ng/ml Mouse TPO (Peprotech), 10 ng/ml Mouse SCF (Peprotech), 10 µM HEPES (Gibco), 1 × Insulin-Transferrin-Selenium-Ethanolamine (ITS-X) (Gibco) and 1 × penicillin–streptomycin (P/S) (Gibco) in Fibronectin-coated plates (Corning). Cells were cultured at 37°C in a 5% CO_2_, atmosphere, and medium was changed every 2 days.

### Genome editing

To prepare the Cas9/RNP, tracrRNA and target-specific crRNA (both from IDT) were mixed, heated at 95°C for 5 minutes, then cooled to room temperature to promote the formation of the RNA duplex. The resulting duplex was then combined with Cas9 protein (IDT) to a final concentration of 18.6 μM, then incubated at room temperature for 20 minutes to assemble the Cas9/RNP^27^.

AAV vectors were used as donor DNA for genome editing. The recombinant expression cassette was packaged into AAV capsids by triple-plasmid transfection of AAVpro293T cells (Takara Bio). AAV vectors were purified from the transfected cells by ultracentrifugation, as described previously^28,29^. The titer of AAV vectors was determined by quantitative PCR (qPCR) of the SV40 polyadenylation signal. The primers and probe used for qPCR were: Fw: GCAATAGCATCACAAATTTCAC; Rv: GATCCAGACATGATAAGATACATTG; Probe: TCACTGCATTCTAGTTGTGGTTTGTCCA.

For delivery of the Cas9/RNP (final concentration: 0.744 µM), the Lonza 4D-Nucleofector™ system was used. Electroporation programs were selected as follows: CM-137 for mouse LK cells, 32D cells, and hairy and enhancer of split 1 (Hes1)-transduced mouse common myeloid progenitors (Hes1-CMP)^30^. Immediately after nucleofection, mouse cells were transduced with AAV donor DNA at 1 × 10⁴ viral genomes (vg)/cell. Culture medium was replaced 24 hours post-nucleofection to remove residual AAV particles. To enhance KI efficiency, ATM inhibitors were added to the culture medium immediately after genome editing. For vehicle control, DMSO at a final concentration matched to the ATM inhibitor condition, 0.0001% v/v was added. The medium was changed after 24 hours to remove compound. A list of ATM inhibitors used is provided in Supplemental Table 1. Following treatment, cells were cultured for an additional 2–5 days before analysis.

For mEGFP KI experiments, KI efficiency was assessed by flow cytometry (Sony SH800) based on the percentage of mEGFP⁺ cells among total live cells or specific fraction such as CD201^+^ CD150^+^ cKit^+^ Sca1^+^ Lineage^-^ for cultured HSCs. All antibodies used in flow cytometry are listed in Supplemental Table 2.

For X-SCID mouse treatment experiments, KI efficiency for Interleukin-2 receptor subunit gamma (*Il2rg*) correction was evaluated by Sanger sequencing of PCR amplicons derived from genome-edited cells. Sequencing traces were analyzed using the ICE CRISPR Analysis online tool (v3.0, 2025; EditCo Bio) to quantify editing outcomes. The primers used for determining KI and KI efficiency are: F1: ACCAGACATTTGTTGTCCAG; F2: TTCCTAGAGCCCCTGAGAACC; R1: CTGACGTGGCAGAACCGT.

### Serial bone marrow transplantation in mice

The genome edited 2 × 10^5^ LK cells were cultured for 3 days and transplanted into 9.5 Gy irradiated C57BL/6J (CD45.2) mice through the retro-orbital vein injection. As accessory cells, 1 × 10^5^ C57BL/6J (CD45.2) bone marrow cells were used. 12 weeks post-transplantation, 5 × 10^6^ total bone marrow cells from primary recipient was transplanted to secondary recipient C57BL/6J (CD45.2) mice post 9.5 Gy irradiation.

Peripheral blood was analyzed at 8 weeks post-transplantation. Around 100 µl of peripheral blood (PB) from recipient mice were collected through submandibular vein in ETDA collection tube. Complete blood count analysis was done using Celltac α automated hematology analyzer. For flow cytometry analysis, the collected PB was centrifuged to separate plasma. Then erythrocytes were lysed by 1 × RBC lysis buffer (Pluriselect). Then washed WBC were used for flow cytometry analysis (BD LSRFortessa™ X-20).

Bone marrow cells were analyzed at 12 weeks post-transplantation. The bone marrow cells were harvested from recipient mice with above mentioned method. After erythrocyte lysis, Lineage^-^ cells were removed by AutoMACS with Depletes mode. These enriched cells were used for flow cytometry analysis. CD150^+^ CD48^-^ cKit^+^ Sca1^+^ Lineage^-^ cells were evaluated as HSC fraction.

### Phosphoproteomics analysis

LK cells were cultured *ex vivo* for 7 days and genome-edited using Cas9/RNP and AAV donor DNA. Cells were collected at 2, 6, or 24 hours post-nucleofection, washed three times with ice-cold PBS, centrifuged to form a cell pellet, snap-frozen in liquid nitrogen, and stored at −80°C until further processing. Phosphoproteomic analysis, including sample preparation, phosphopeptide enrichment, LC-MS/MS acquisition, and data processing, was outsourced to BGI Genomics and performed according to the provider’s standard protocols. Peptide identification was carried out using Proteome Discoverer 1.4 (Thermo Fisher Scientific) integrated with MASCOT 2.3 for database searching. Percolator was used to reprocess the Mascot search results, and peptides were filtered using a false discovery rate (FDR) threshold of ≤ 1.0% at the spectral level. Phosphorylation site localization was performed using the phosphoRS module in Proteome Discoverer, and phosphosites were considered confident when the phosphoRS probability was ≥ 0.75. Functional annotation and motif enrichment analyses were conducted for proteins containing confidently localized phosphosites.

### Capillary western blotting

LK cells were cultured *ex vivo* for 7 days and genome-edited using Cas9/RNP and AAV donor DNA. Cells were collected at 3-4 hours post-nucleofection, washed with ice-cold PBS, and centrifuged to form a cell pellet. The pellet was lysed using M-PER (Thermo Fisher Scientific) containing protease inhibitors (Nacalai Tesque) and a proteinase inhibitor cocktail (Abcam). The extracted proteins were mixed with 2 × Laemmli buffer (Bio-Rad) containing dithiothreitol at a final concentration of 100 mM and boiled at 95°C for 5 minutes. Then, protein concentration was determined using the Pierce™ 660 nm Protein Assay (Thermo Fisher Scientific). Capillary western blotting was performed using the JESS system (Bio-techne). For ATM and p-ATM, the 66-440 kDa Fluorescence Separation Module was used; for GAPDH, NPM1, p-NPM1, p53, p-p53, CHK2 and p-CHK2, the 12-230 kDa Fluorescence Separation Module was used; and for p-H2AX, p21, Caspase 3 and Cleaved Caspase 3, the 2-40kDa Fluorescence Separation Module was used, all according to the manufacturer’s protocol. A list of antibodies used is provided in Supplemental Table 3.

### Flow cytometry with Cellview

7-day cultured LK cells were genome edited with Cas9/RNP and AAV donor DNA. Cells were collected at 3 hours post-nucleofection and washed with PBS and stained with antibodies for 30 minutes at 4°C. Surface markers were stained using antibodies against Lineage markers (CD3, B220, Gr1, CD11b, Ter119), c-Kit, Sca-1, to define hematopoietic stem and progenitor cell populations. For intracellular staining (γH2AX), cells were fixed and permeabilized using the BD Phosflow™ Fix Buffer I and BD Phosflow™ Perm Buffer III (BD Biosciences). Samples were acquired on a BD FACSDiscover™ S8 Cell Sorter with BD CellView™ Image Technology and BD SpectralFX™ Technology (BD Biosciences). Gates were defined based on fluorescence minus one or isotype controls where applicable. Data were analyzed using FlowJo™ 10 software.

### Statistical analysis

Statistical analyses were performed using GraphPad Prism 10. Data are presented as mean ± standard deviation. Percentage values were arcsine-transformed prior to analysis, and normality was assessed using the Shapiro–Wilk test. Depending on distribution, parametric or nonparametric tests were applied, including unpaired t-tests, Mann–Whitney U tests, Tukey’s multiple comparison tests and Holm–Šidák multiple comparison tests. All tests were two-sided with a 95% confidence interval, and p-values < 0.05 were considered statistically significant. Experiments were conducted in biological triplicate unless otherwise indicated; transplantation assays used six mice per group. Specific statistical tests are detailed in the figure legends.

## Results

### ATM inhibition enhances KI efficiency in hematopoietic cell lines

To test whether ATM inhibition on enhancement of KI efficiency in hematopoietic cells, we used fluorescent-based KI reporter system. To explain, a promoterless monomeric EGFP (mEGFP) coding sequence was inserted into exon 6 of mouse *beta Actin* (*Actb)* gene using a donor DNA flanked by 1-kilobase (kb) 5’ and 3’ homology arms (HA) (Figure 1A). Because the donor DNA construct lacks an exogenous promoter, mEGFP expression occurs only when the allele is precisely genome-edited via HDR, allowing accurate quantification of KI efficiency. (Figure 1B).

**Figure 1.**
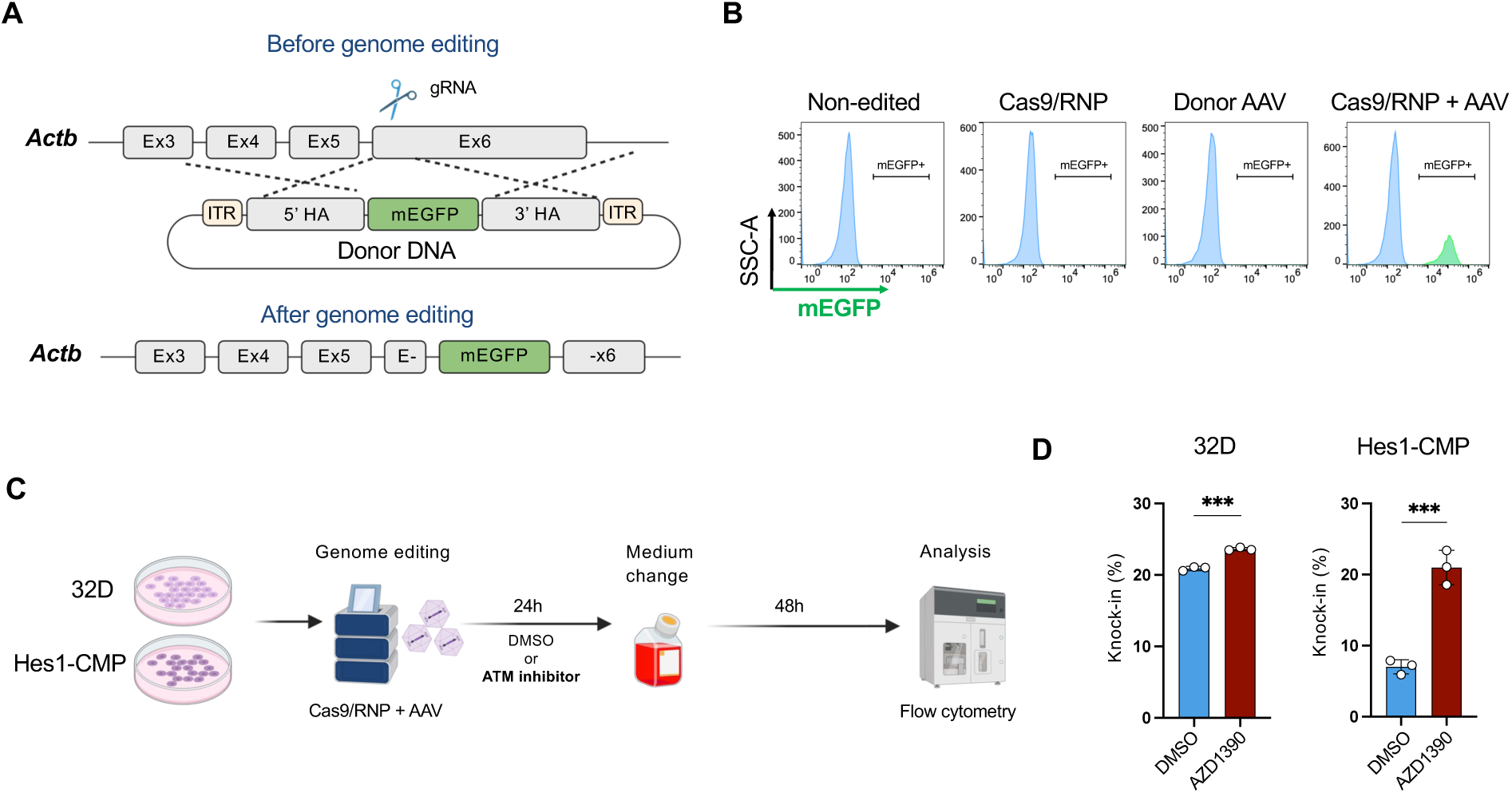
ATM inhibition enhances knock-in (KI) efficiency in cell lines. (A) Diagram of the mEGFP KI construct. (B) Representative flow cytometry results. When mEGFP is knocked-in into *Actb* in 32D cells, the mEGFP expression is detected by flow cytometry. (C) Schematic of the experimental procedure for genome editing in cell lines. (D) KI efficiency following 10 nM AZD1390 treatment in 32D cells and Hes1-CMP cells. Data are presented as the mean ± standard deviation (n=3). Unpaired t-tests were conducted. For 32D cells, *P* = 0.0002; for Hes1-CMP cells *P* = 0.0005.

To assess whether ATM inhibition could enhance KI efficiency in hematopoietic cells, we tested two mouse hematopoietic cell lines: 32D Clone 3 cell line, and Hes1-CMP cells^30^. Cells were electroporated with Cas9/RNP, transduced with AAV donor DNA, and treated with or without 10 nM AZD1390, a highly potent and selective ATM inhibitor, for 24 hours (Figure 1C). Flow cytometric analysis revealed that AZD1390 treatment significantly increased the percentage of mEGFP⁺ cells in both cell lines compared to DMSO controls (Figure 1D). These results indicate that transient ATM inhibition enhances KI efficiency in hematopoietic cell lines.

### ATM inhibition increases KI efficiency in *ex vivo* cultured primary mouse HSCs

We next evaluated the effect of ATM inhibition on genome editing in primary mouse HSCs. Lineage^-^ cKit^+^ (LK) cells were isolated from mouse bone marrow (BM) and cultured for 7 days *ex vivo* using HemEx-Type9A media which contains polyvinyl alcohol to promote HSCs expansion^31,32^ (Figure 2A). The same KI reporter described above was used to detect and quantify KI by the mEGFP-expressing cells. KI efficiency was assessed in total cultured LK cells or in phenotypically defined HSC population (CD201^+^ CD150^+^ c-Kit^+^ Sca1^+^ Lineage^-^; referred to as CD201^+^ CD150^+^ KSL) using flow cytometry (Figure 2A-B; Supplemental Figure 1). To evaluate the effect of ATM inhibition, we tested several ATM inhibitors: AZD1390, M4076, KU60019, AZ32, and KU55933. All compounds showed a tendency to increase KI efficiency in both total cells and in the HSC fraction. Among them, 10 nM AZD1390 exhibited the most significant effect, increasing KI efficiency from 43.0 ± 0.7% to 62.2% ± 0.4 in total cells, and from 44.4% ± 6.4% to 70.7% ± 6.6% in the HSC fraction (*P* < 0.0001 and *P* < 0.0082, respectively). In addition, quantification of mEGFP⁺ HSCs revealed that ATM inhibitor treatment significantly enhanced KI efficiency within the HSC population. (Figure 2C-D, Supplemental Figure 2). This effect was observed in a dose-dependent manner with AZD1390 (Supplemental Figure 3). These results suggest the transient ATM inhibition significantly enhances KI efficiency in *ex vivo*-cultured primary mouse HSCs.

**Figure 2.**
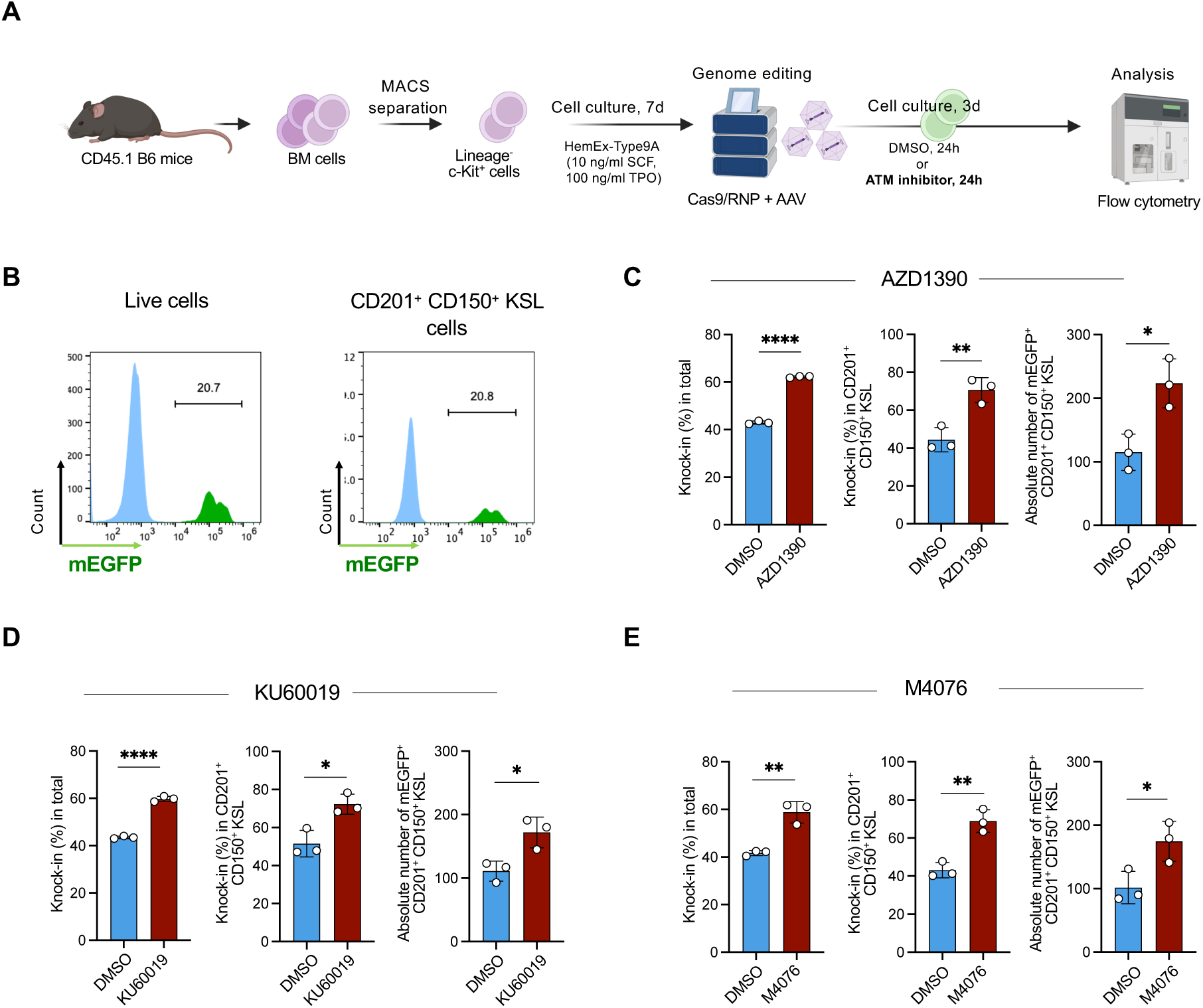
ATM inhibition enhances knock-in (KI) efficiency in mouse HSCs *ex vivo*. (A) Schematic of the *ex vivo* genome editing protocol in mouse HSCs. (B) Representative flow cytometry plots. KI efficiency was assessed in live cells and CD201^+^ CD150^+^ c-Kit^+^ Sca1^+^ Lin^-^ cells (HSCs). (C) Genome editing results following treatment with 10 nM AZD1390. Data are presented as the mean ± standard deviation (SD) (n=3). KI (%) in total, Unpaired t-tests were conducted. *P* < 0.0001; For KI (%) in HSCs, *P* = 0.0082; For Absolute number of mEGFP^+^ HSCs, Unpaired t-tests were conducted. *P* = 0.0173. (D) Genome editing outcomes following treatment with 1 µM KU60019. Data are presented as the mean ± SD (n=3). For KI (%) in total, Unpaired t-tests were conducted. *P* < 0.0001; For KI (%) in HSCs, Unpaired t-tests were conducted. *P* = 0.0154; For Absolute number of mEGFP^+^ HSCs, Unpaired t-tests were conducted. *P* = 0.0214. (E) Genome editing outcomes following treatment with 1 µM M4076. Data are presented as the mean ± SD (n=3). For KI (%) in total, Unpaired t-tests were conducted. *P* = 0.0032; For KI (%) in HSCs, Unpaired t-test were conducted. *P* = 0.0039; For Absolute number of mEGFP^+^ HSCs, *P* = 0.0361.

### HSCs genome-edited with ATM inhibitors retain long-term, multilineage repopulating capacity

To determine whether ATM inhibition preserves the long-term repopulating capacity of genome-edited HSCs, we performed serial transplantation using CD45.2^+^ recipient mice and CD45.1^+^ donor mice, both on C57BL/6 background (Figure 3A). LK cells harvested from donor mouse BM, were cultured *ex vivo* for 7 days with following genome-edited using Cas9/RNP and AAV donor DNA, with or without 10 nM AZD1390, and transplanted into lethally irradiated recipient mice (n=6 per group).

**Figure 3.**
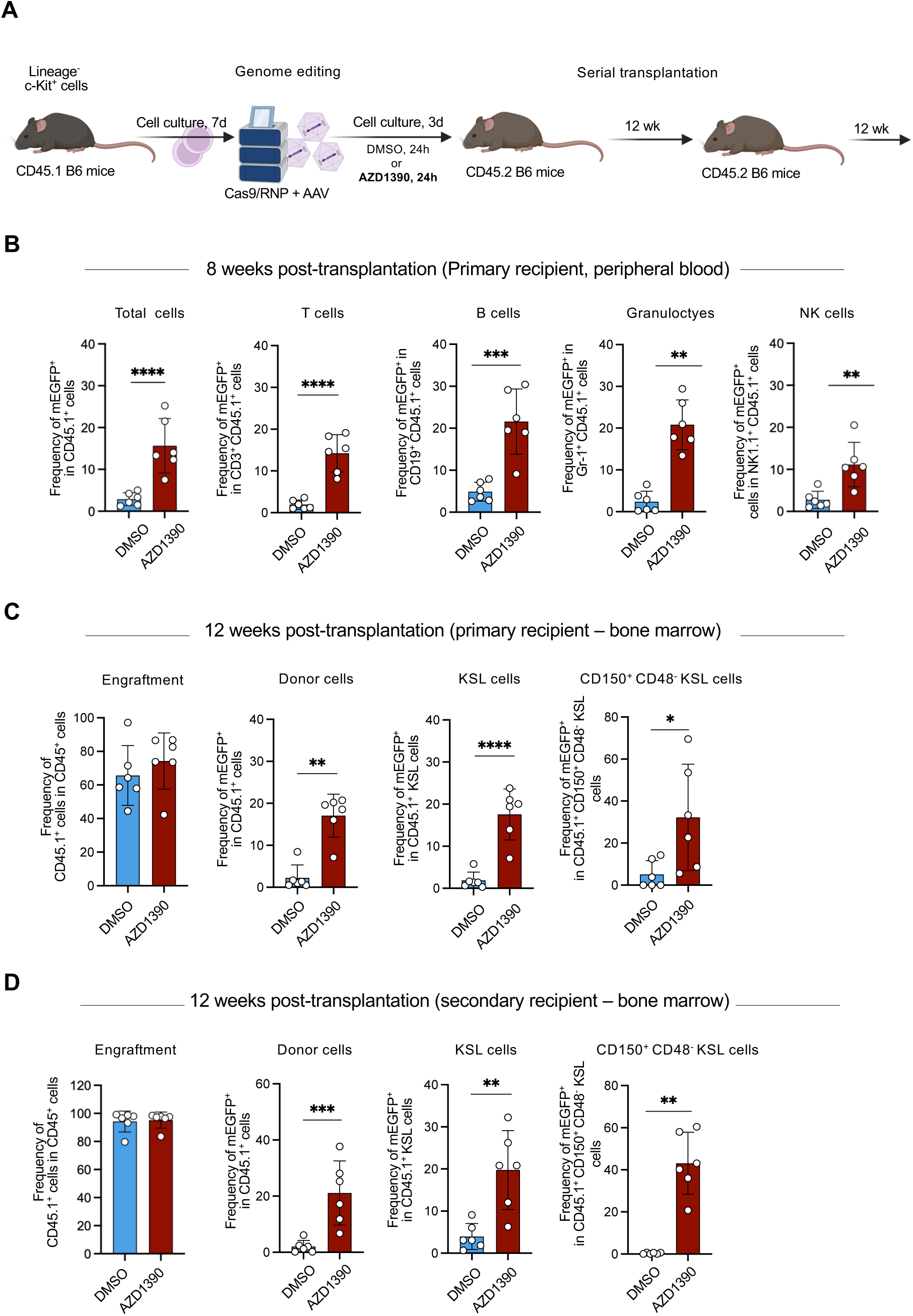
ATM inhibition enhances knock-in (KI) efficiency in functional HSCs. (A) Schematic of the experimental procedure for the transplantation analysis. (B) Peripheral blood (PB) analysis at 8 weeks post-transplantation (primary). Data are shown as the mean ± standard deviation (SD) (n=6). For frequency of mEGFP^+^ cells in donor cells, Unpaired t-test was conducted. *P* < 0.0001; For frequency of mEGFP^+^ cells in donor T cells, Unpaired t-test was conducted. *P* < 0.0001; For frequency of mEGFP^+^ cells in donor B cells, Unpaired t-test was conducted. *P* = 0.0002; For frequency of mEGFP^+^ cells in donor granulocytes, Mann-Whitney t-test was conducted. *P* = 0.0022; For frequency of mEGFP^+^ cells in donor NK cells, Unpaired t-test was conducted. *P* = 0.0017. (C) Bone marrow (BM) analysis at 12 weeks post-transplantation (primary). Data are shown as the mean ± SD (n=6). For engraftment rate, Unpaired t-test was conducted. *P* = 0.502; For frequency of mEGFP^+^ cells in donor cells, Mann-Whitney t-test was conducted. *P* = 0.0043; For frequency of mEGFP^+^ cells in donor KSL cells, Unpaired t-test was conducted. *P* < 0.0001; For frequency of mEGFP^+^ cells in donor CD150^+^ CD48^-^ KSL cells (HSCs), Unpaired t-test was conducted. *P* = 0.014. (D) BM analysis at 12 weeks post-transplantation (secondary). Data are shown as the mean ± SD (n=6). For engraftment rate, Mann-Whitney t-test was conducted. *P* = 0.8182; For frequency of mEGFP^+^ cells in donor cells, Unpaired t-test was conducted. *P* = 0.0005; For frequency of mEGFP^+^ cells in donor c-Kit^+^ Sca1^+^ Lin^-^ cells, Unpaired t-test was conducted. *P* = 0.0016; For frequency of mEGFP^+^ cells in donor HSCs, Mann-Whitney t-test was conducted. *P* = 0.0022.

To assess hematopoietic reconstitution, we performed Complete Blood Count (CBC) analysis at 4 weeks post-transplantation. No significant differences were observed between DMSO and AZD1390-treated groups in red blood cell count, white blood cell (WBC) count, platelet count, hematocrit and hemoglobin levels (Supplemental Figure 4A). In addition, we analyzed major leukocyte fractions including T cells, B cells, NK cells and granulocytes, in PB at 4- and 8-weeks post-transplantation by flow cytometry. No significant differences were observed between DMSO and AZD1390 groups (Supplemental Figure 4B-C). In both transplantations, CD45.2^+^ whole BM cells were used as accessory cells. To specifically assess KI efficiency in donor-derived cells, we analyzed in CD45.1^+^ compartment. Notably, frequency of mEGFP^+^ cells within CD45.1^+^ cells was significantly higher in AZD1390-treated mice across all major lineages, including T cells, B cells, NK cells and granulocytes compared to DMSO controls (Figure 3B; Supplemental Figure 5A-B). BM analysis at 12 weeks post-transplantation showed significantly increased frequency of mEGFP^+^ cells in AZD1390-treated mice within total CD45.1^+^ cells, CD45.1^+^ KSL cells, and CD45.1^+^ CD150^+^ CD48^-^ KSL cells (HSC fraction). Especially in CD45.1^+^ HSCs, the frequency of mEGFP^+^ cells was 5.1% ± 6.6 % in the DMSO group and 32.4% ± 25.3% in the AZD1390 group (*P* < 0.014) (Figure 3C, Supplemental Figure 6). This elevated frequency of mEGFP⁺ cells in the AZD1390 group was maintained in secondary recipients at 12 weeks post-transplantation, reaching 43.1% ± 14.7%. In contrast, the frequency of mEGFP⁺ HSCs in the DMSO group decreased to 0.33% ± 0.37% (Figure 3D). In addition, serial bone marrow analysis at 12 weeks in both primary and secondary recipients demonstrated comparable engraftment rates between DMSO and AZD1390-treated groups (Figure 3C-D). These results indicate that transient ATM inhibition enhances KI efficiency in functional HSCs without impairing long-term hematopoietic reconstitution.

### ATM inhibition suppresses the overactivation of ATM induced by Cas9/RNP and AAV donor DNA, thereby attenuating downstream p53–Caspase-3 activation

To investigate the mechanism by which ATM inhibition increases KI efficiency in HSCs, we performed phosphoproteomic analysis. 7-day cultured LK cells were genome-edited and treated with DMSO or 10 nM AZD1390. Then samples were prepared for phosphoproteomics analysis at 2 hours, 6 hours and 24 hours post-genome editing (Figure 4A). Principal component analysis of phosphopeptide signatures showed the clear separation between the DMSO and AZD1390 groups in the 2-hour time-point, indicating ATM’s distinct signaling profiles after inhibition (Figure 4B). Phosphopeptide analysis revealed that p53 regulators shifted from an activated state in the DMSO group to a suppressed state in the AZD1390 group at 2 hours and 6 hours post-genome editing. Interestingly, Nucleophosmin 1 (NPM1), a multifunctional nucleolar protein involved in both p53 regulation and genomic stability in hematopoietic cells, exhibited increased phosphorylation at Ser10 in the AZD1390 group. (Figure 4C). We then validated these findings by performing capillary western blotting at 3–4 hours post-genome editing, between the 2- and 6-hour time points (Figure 4D). From this analysis, cells in the DMSO group exhibited overactivation of ATM at Ser1987, which was significantly suppressed by AZD1390 treatment. In addition, phosphorylated NPM1 at Ser10 was significantly increased in the AZD1390 group, a modification crucial for genomic stability (Figure 4F). Then we tested one of key molecules for DDR and ATM’s downstream; phospho-H2AX, so-called γ-H2AX by flow cytometry with CellView^TM^ (Supplemental Figure 7). As expected, the number of γ-H2AX foci was higher in the DMSO group, whereas AZD1390 treatment markedly reduced the formation of these foci (Figure 4G-H). These results were further supported by capillary western blot analysis. In the DMSO group, we observed activation of p53 and subsequent activation of Caspase-3, both of which were suppressed by ATM inhibition (Figure 4I-J). These findings indicate that ATM inhibition suppresses Cas9-AAV-mediated DDR and subsequent p53-dependent apoptosis (Figure 4K).

**Figure 4.**
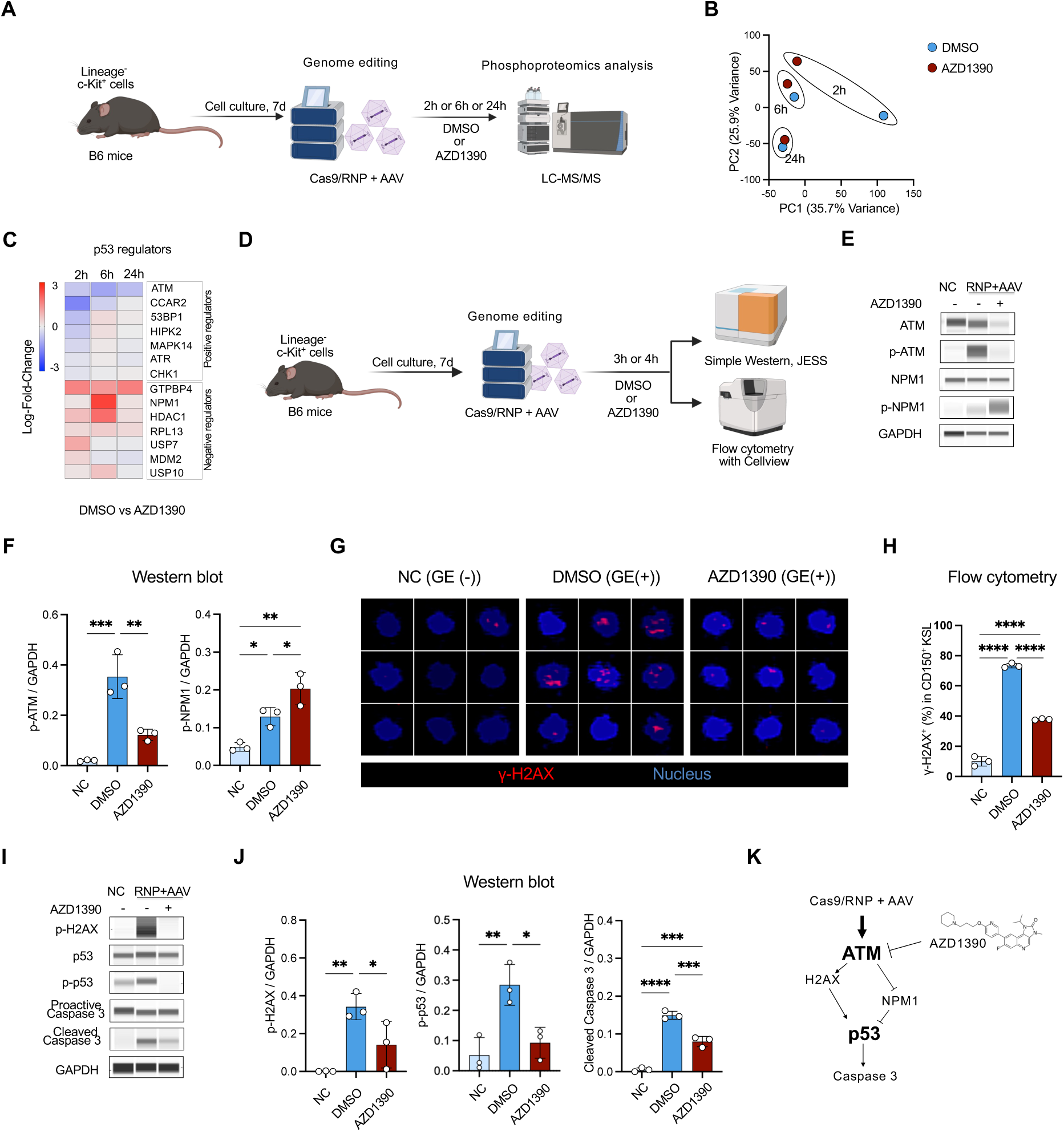
ATM inhibition suppresses Cas9-AAV-induced DNA damage response (DDR) and subsequent p53-dependent apoptosis. (A) Schematic of the experimental procedure for phosphoproteomic analysis. (B) Principal component analysis (PCA) results of the phosphopeptides after genome editing with Cas9/RNP and AAV donor DNA in the presence or absence of ATM inhibitor at 2-, 6- and 24-hours post-genome editing. (C) Heatmap showing changes of phosphorylation level of p53 regulators at 2-, 6- and 24-hours post-genome editing, comparing DMSO and AZD1390-treated groups. Log-transformed fold changes were used. (D) Schematic of the experimental procedure for capillary western blotting and flow cytometry analysis. (E) Western blotting results showing genome editing with Cas9/RNP and AAV donor DNA induces ATM activation (Ser-1987) at 4 hours post-genome editing. 10 nM AZD1390 treatment suppresses the activation of ATM and also enhances the activation of NPM1 (Ser-10). (F) Quantification of the western blot results. The signal intensities were normalized to GAPDH. Data are presented as the mean ± standard deviation (SD) (n=3). For p-ATM, Tukey’s multiple comparisons tests were conducted. NC vs DMSO, *P* = 0.0005; NC vs AZD1390, *P* = 0.1144; DMSO vs AZD1390, *P* = 0.0038. For p-NPM1, Tukey’s multiple comparisons tests were conducted. NC vs DMSO, *P* = 0.035; NC vs AZD1390, *P* = 0.0016; DMSO vs AZD1390, *P* = 0.0477. (G) Representative images of flow cytometry analysis with Cellview^TM^. The AZD1390 treatment decreases the γ-H2AX^+^ cells compared to DMSO control. GE stands for genome-editing with Cas9/RNP and AAV donor DNA. (H) Flow cytometry quantification of the γ-H2AX activation analysis. Data are presented as the mean ± SD (n=3). Tukey’s multiple comparisons tests were conducted. NC vs DMSO, *P* < 0.0001; NC vs AZD1390, *P* < 0.0001; DMSO vs AZD1390, *P* < 0.0001. (I) Western blotting results showing Cas9/RNP and AAV donor DNA genome editing activates H2AX (Ser-139), p53 (Ser-15) and Caspase 3. 10 nM AZD1390 treatment suppresses the activation of H2AX, p53 and following Caspase 3. (J) Quantification of the western blot results. The signal intensities were normalized to GAPDH. Data are presented as the mean ± SD (n=3). For p-H2AX, Holm-Šídák’s multiple comparisons tests were conducted. NC vs DMSO, *P* = 0.0065; NC vs AZD1390, *P* = 0.0799; DMSO vs AZD1390, *P* = 0.0468. For p-p53, Tukey’s multiple comparisons tests were conducted. NC vs DMSO, *P* = 0.0073; NC vs AZD1390, *P* = 0.6937; DMSO vs AZD1390, *P* = 0.0179. For Cleaved Caspase 3, Tukey’s multiple comparisons tests were conducted. NC vs DMSO, *P* < 0.0001; NC vs AZD1390, *P* = 0.0003; DMSO vs AZD1390, *P* = 0.0005. (K) Schematic explanation of Cas9-AAV-induced ATM overactivation and subsequent p-53 dependent apoptosis, which is mitigated by transient ATM inhibition.

### ATM inhibition enhances the efficiency of therapeutic genome editing to correct the *Il2rg* mutation in the X-SCID mouse model

We next evaluated whether our optimized genome editing approach, in combination with ATM inhibition, could effectively correct the *Interleukin 2 receptor subunit gamma* (*Il2rg)* mutation responsible for X-SCID. For this purpose, we used a previously established X-SCID mouse model carrying a single-nucleotide insertion in exon 4 of *Il2rg*, which causes a frameshift mutation and loss of IL2RG (CD132) function^15^. To repair the mutation, we designed a gene correction strategy using AAV donor DNA containing silent mutations in DSB site, flanked by 2 kb HAs at both the 5’ and 3’ homology ends (Figure 5A). Precise knock-in restores the native amino acid sequence and functional protein expression (Figure 5B). Before applying this approach to hematopoietic cells, we first validated the genome editing system in bone marrow–derived stromal cells immortalized with SV40 large T antigen. These cells were genome-edited with Cas9/RNP and AAV donor DNA and then cultured for 5 days. PCR amplification using a primer pair spanning the 5’ HA (F1) and a downstream region outside the 3’ HA (R1) was followed by Sanger sequencing. KI efficiency was quantified using ICE CRISPR Analysis (Figure 5C, Supplemental Figure 8). Treatment with 10 nM AZD1390 significantly increased KI frequency compared to DMSO control (Figure 5D). We then applied the same genome editing protocol to 7-day ex vivo–cultured LK cells from X-SCID mice. To confirm KI in *Il2rg* mutation site, we used a primer pair (F2–R1) designed to incorporate only in the region of “silent mutations” and downstream genomic sequence. Gel electrophoresis of PCR amplicons showed successful KI in LK cells (Figure 5E). To quantify KI efficiency, we employed the same approach used for stromal cells. Sanger sequencing and ICE analysis revealed significantly enhanced KI efficiency in the AZD1390 group (Figure 5F). Finally, we assessed expression of CD132 in KSL cells by flow cytometry. While CD132 expression is typically low in KSLs, AZD1390-treated cells exhibited slightly higher CD132 expression levels compared to the DMSO group and increased frequency of CD132^+^ cells (Figure 5G), indicating successful functional correction of the *Il2rg* mutation.

**Figure 5.**
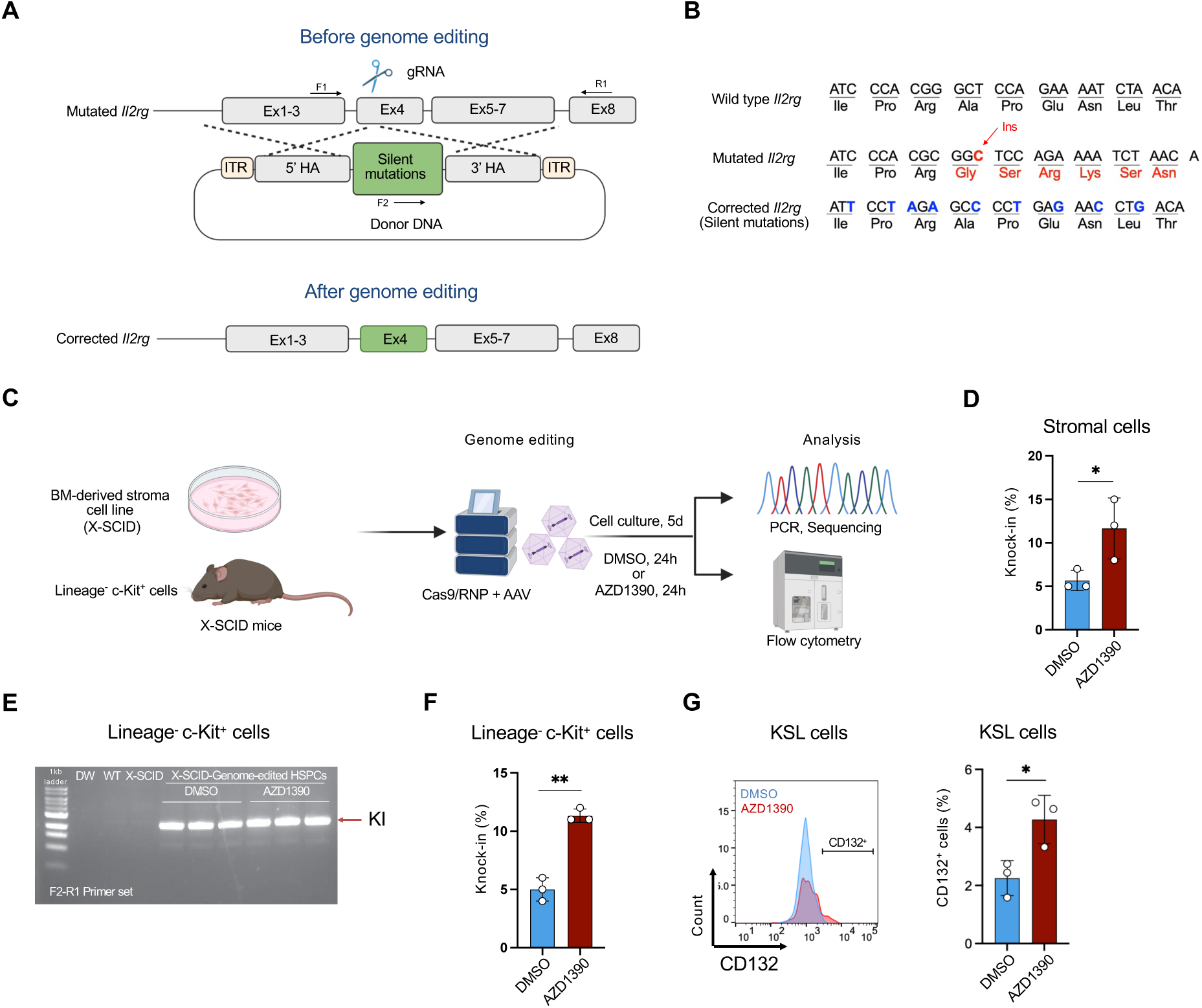
ATM inhibition enhances knock-in (KI) efficiency in X-SCID mouse-derived HSPCs. (A) Diagram of the genome editing construct for X-SCID treatment. (B) Illustration of “silent mutation” strategy used to repair the *Il2rg* mutation. (C) Schematic of the experimental procedure for gene correction targeting the *Il2rg* mutation. (D) KI efficiency in bone marrow-derived stromal cells treated with 10 nM AZD1390 treatment. Data are presented as the mean ± standard deviation (SD) (n=3). Unpaired t-tests were conducted. *P* = 0.039. (E) Gel electrophoresis results demonstrating successful KI in X-SCID mouse bone marrow-derived HSPCs. (F) KI efficiency following 10 nM AZD1390 treatment in Lineage^-^ c-Kit^+^ cells. Data are presented as the mean ± SD (n=3). Unpaired t-tests were conducted. *P* = 0.0012. (G) Representative flow cytometry histograms showing CD132 (IL2RG) expression in DMSO and AZD1390-treated groups. Frequency of CD132^+^ cells following treatment of 10 nM AZD1390. Data are presented as the mean ± SD (n=3). Unpaired t-tests were conducted. *P* = 0.0009.

## Discussion

In this study, we demonstrate that transient inhibition of ATM enhances KI efficiency in mouse (CD201⁺ CD150⁺ KSL *ex vivo* and CD150⁺ CD48⁻ KSL *in vivo*) HSCs. This improvement is achieved by suppressing the DDR triggered by Cas9 and AAV donor delivery, which otherwise induces excessive ATM activation and subsequent p53-dependent apoptosis.

ATM is a key regulator of the DDR and is one of the earliest sensors of DSB. Under conditions such as ionizing radiation (IR) or treatment with genotoxic agents like etoposide, multiple DSBs activate ATM, leading to apoptosis via p53 and caspase-3 signaling^24,33,34^. During genome editing, however, CRISPR/Cas9 typically generates targeted DSB^35^. Early ATM activation leads to CHK2–p53–p21 signaling, inducing G1 arrest that favors NHEJ^36^. If the DSB is not repaired by NHEJ, HDR may occur, particularly during the S and G2 phases of the cell cycle. Importantly, even a single unrepaired DSB can activate apoptotic pathway^37^. Notably, in genome editing using Cas9/RNP and AAV donor DNA, high AAV copy number appears to mimic multiple DSBs, leading to ATM overactivation and apoptosis. While a higher amount of donor DNA increases the likelihood of successful KI, it paradoxically induces cell death in KI-competent cells. Our findings demonstrate that ATM inhibition mitigates this detrimental overactivation, thereby preserving cells with high KI potential.

In addition, phosphoproteomic analysis revealed that ATM inhibition modulates early DDR signaling. Specifically, phosphorylation of canonical ATM substrates was reduced, whereas phosphorylation of NPM1 at Ser10 was increased. NPM1 is a multifunctional protein involved in maintaining genomic stability, regulating chromatin structure, and facilitating DNA repair^38^. While NPM1 mutations in acute myeloid leukemia impair p53 signaling^39^, wild-type NPM1 can promote cell cycle progression by regulating CDK1. Phosphorylation of NPM1 at Ser10 is associated with G2/M transition and chromatin condensation, cellular processes that are favorable for HDR^40^. Furthermore, ATM inhibition led to decreased CHK2 phosphorylation and reduced p21 protein levels (Supplemental Figure 10), which may relieve inhibition of CDK1 and thereby facilitate entry into S/G2 phase, which are conductive to KI^41^. Although we did not directly assess cell cycle dynamics in this study, our data suggest that ATM inhibition may enhance KI efficiency by enhancing transition into cell cycle phases optimal for HDR.

Our primary goal was to enhance KI-mediated genome editing in HSCs, which are intrinsically quiescent and typically favor NHEJ. To address this, we used *ex vivo* culture to stimulate mouse HSC cycling and promote KI^42^. After 7-day culture of mouse HSPCs, we obtained ∼40% KI efficiency using Cas9/RNP and AAV donor DNA. However, without ATM inhibition, transplantation experiments showed rapid loss of genome-edited HSCs post-engraftment, and only ∼0.3% of genome-edited HSCs were detected in the bone marrow 12 weeks after secondary transplantation. This rapid loss of genome-edited HSCs is most likely attributable to sustained activation of the DDR pathway and subsequent apoptosis^43^. In previous report, AAV transduction alone in human CD34^+^ HSPC induces massive phosphorylation of H2AX that is another key molecule for DDR^44^. In addition, AAV delivery of donor DNA activates the p53-related pathway and induces apoptosis in human mobilized PB HSPCs^45^, and transient expression of dominant negative p53 protein during genome editing increased HDR efficiency^46^. In our study, transient ATM inhibition reduced phosphorylation of H2AX and suppressed p53 activation, thereby attenuating the DDR. In transplantation experiment, over 40% of genome-edited HSCs were preserved at 12 weeks post-secondary transplantation in AZD1390 group. These findings suggest that ATM inhibition not only enhances KI efficiency but also preserves long-term repopulating HSCs by preventing stress-induced apoptosis.

To assess the therapeutic relevance of our approach, we applied it to a mouse model of X-SCID caused by a frameshift mutation in *Il2rg*. Because corrected lymphoid cells gain a strong selective advantage, even modest KI efficiency can restore immune function^47^. Based on results from mEGFP KI experiment, ∼0.3% of KI in HSCs might not be enough to cure X-SCID mice, therefore ATM inhibition may be needed for X-SCID mouse treatment. Our findings show a significant increase of KI efficiency for X-SCID treatment, and restoration of CD132 expression, however, demonstrating the successful KI in functional HSCs will require transplantation experiments, which remains a subject for further investigation.

In conclusion, our study demonstrates that transient ATM inhibition strikingly enhances KI efficiency, preserves the long-term repopulating capacity of genome-edited HSCs, and enables effective functional correction in disease models. This approach provides a promising strategy and strong foundation for advancing genome editing toward safer and more efficient therapeutic applications and may help pave the way for future clinical translation.

## Acknowledgements

We thank Hiroko Hayakawa for her technical assistance with flow cytometry analysis, Takeshi Tokuyama for his support with capillary western blotting, and Mika Kishimoto for assistance with AAV production. We also thank Tomoya Abe for his help with phosphoproteomics analysis. In addition, we acknowledge Hideki Uosaki for his constructive suggestions. Some part of figures was created with Biorender.com.

This study was supported by the Japan Agency for Medical Research and Development (AMED) [JP23bm1123020 to F.N.].

## Authorship

### Contribution

M.N., H.H. and K.I. performed experiments and analyzed data; Y.K. and T.O. produced AAV vectors; H.H., Y.H. and F.N. conceived of the study; F.N. supervised the study; M.N. and F.N. interpreted the data and wrote the manuscript; all authors critically revised and approved the final version.

### Conflict-of-interest disclosure

H.H. and Y.H. were affiliated with a joint laboratory between Jichi Medical University and Sumitomo Pharma Co., Ltd. The remaining authors declare no competing financial interests.

Correspondence: Fumio Nakahara, Division of Regenerative Medicine, Center for Molecular Medicine, Jichi Medical University, Shimotsuke, Tochigi, Japan;

