## Supplemental data for "Transient ATM inhibition enhances knock-in efficiency in hematopoietic stem cells by attenuating the DNA damage response"

**Supplemental Table 1.** List of the ATM inhibitors

| No | Name | Manufacturer | Product code | Concentration for mouse cells |
|---|---|---|---|---|
| 1 | KU55933 | Selleck | S1092 | 1 μM |
| 2 | KU60019 | Selleck | S1570 | 1 μM |
| 3 | AZD1390 | Selleck | S8680 | 10 nM |
| 4 | AZ32 | Selleck | S8556 | 1 μM |
| 5 | M4076 | Selleck | E1057 | 1 μM |

**Supplemental Table 2.** List of the antibodies for flow cytometry analysis

| No | Name | Manufacturer | Product code |
|---|---|---|---|
| 1 | Alexa Fluor® 700 anti-mouse Lineage Cocktail | Biolegend | 133313 |
| 2 | APC anti-mouse CD48 Antibody | Biolegend | 103412 |
| 3 | APC/Cyanine7 anti-mouse CD45.1 Antibody | Biolegend | 110716 |
| 4 | BD Horizon™ Brilliant Stain Buffer Plus | BD Biosciences | 566385 |
| 6 | BD Pharmingen™ 7-AAD | BD Biosciences | 559925 |
| 8 | BD Pharmingen™ PE Mouse Anti-H2AX (pS139) | BD Biosciences | 562377 |
| 9 | BD Pharmingen™ PE Mouse IgG1, κ Isotype Control | BD Biosciences | 554680 |
| 10 | BD Pharmingen™ PE Rat Anti-Mouse CD132 | BD Biosciences | 554457 |
| 12 | Biotin anti-mouse Lineage Panel | Biolegend | 133307 |
| 13 | Brilliant Violet 421™ anti-mouse CD45.1 Antibody | Biolegend | 110731 |
| 14 | Brilliant Violet 421™ anti-mouse Ly-6A/E (Sca-1) Antibody | Biolegend | 108128 |
| 15 | Brilliant Violet 421™ Streptavidin | Biolegend | 405226 |
| 16 | Brilliant Violet 510™ anti-mouse CD117 (c-kit) Antibody | Biolegend | 135119 |
| 17 | Brilliant Violet 510™ Rat IgG2b, κ Isotype Ctrl Antibody | Biolegend | 400645 |
| 18 | Brilliant Violet 711™ anti-mouse CD3 Antibody | Biolegend | 100241 |
| 19 | Brilliant Violet 711™ anti-mouse Ly-6G/Ly-6C (Gr-1) Antibody | Biolegend | 108443 |
| 20 | Brilliant Violet 785™ anti-mouse CD150 (SLAM) Antibody | Biolegend | 115937 |
| 21 | CD117 (c-Kit) Monoclonal Antibody (2B8), APC | Invitrogen | 17-1171-82 |
| 22 | CD117 (c-Kit) Monoclonal Antibody (2B8), PE | Invitrogen | 12-1171-82 |
| 23 | Cellstain® DAPI solution | Dojindo | 340-07971 |
| 24 | DRAQ5™ | Biolegend | 424101 |
| 25 | PE anti-mouse CD19 Antibody | Biolegend | 115507 |
| 26 | PE anti-mouse CD201 (EPCR) Antibody | Biolegend | 141504 |
| 27 | PE anti-mouse NK-1.1 Antibody | Biolegend | 108707 |
| 28 | PE/Cyanine7 anti-mouse CD150 (SLAM) Antibody | Biolegend | 115914 |
| 29 | PE/Cyanine7 anti-mouse CD45.2 Antibody | Biolegend | 109830 |
| 30 | PE/Dazzle™ 594 anti-mouse CD45.2 Antibody | Biolegend | 109846 |
| 32 | PerCP/Cyanine5.5 anti-mouse Ly-6A/E (Sca-1) Antibody | Biolegend | 108124 |

**Supplemental Table 3.** List of the antibodies for western blotting analysis

| No | Name | Manufacturer | Product code |
|---|---|---|---|
| 1 | Anti-ATM (phospho S1987) | Abcam | ab315019 |
| 2 | ATM (D2E2) Rabbit mAb | Cell signaling | 2873 |
| 3 | Caspase-3 Antibody | Cell signaling | 9662 |
| 4 | Cleaved Caspase-3 (Asp175) Antibody | Cell signaling | 9661 |
| 5 | GAPDH Antibody (6C5) | Santa Cruz | sc-32233 |
| 6 | NPM1 Antibody | Cell signaling | 3542S |
| 7 | p53 (1C12) Mouse mAb | Cell signaling | 2524 |
| 8 | Phospho-Histone H2A.X (Ser139) (20E3) Rabbit mAb | Cell signaling | 9718 |
| 9 | Phospho-NPM1 (Ser4) Polyclonal Antibody | Invitrogen | PA5-106194 |
| 10 | Phospho-p53 (Ser15) Antibody | Cell signaling | 9284 |
| 11 | Chk2 Antibody | Cell signaling | 2662 |
| 12 | Phospho-Chk2 (Thr68) | Affinity Biosciences | AF3036 |
| 13 | p21 Waf1/Cip1 Antibody | Cell signaling | 64016 |

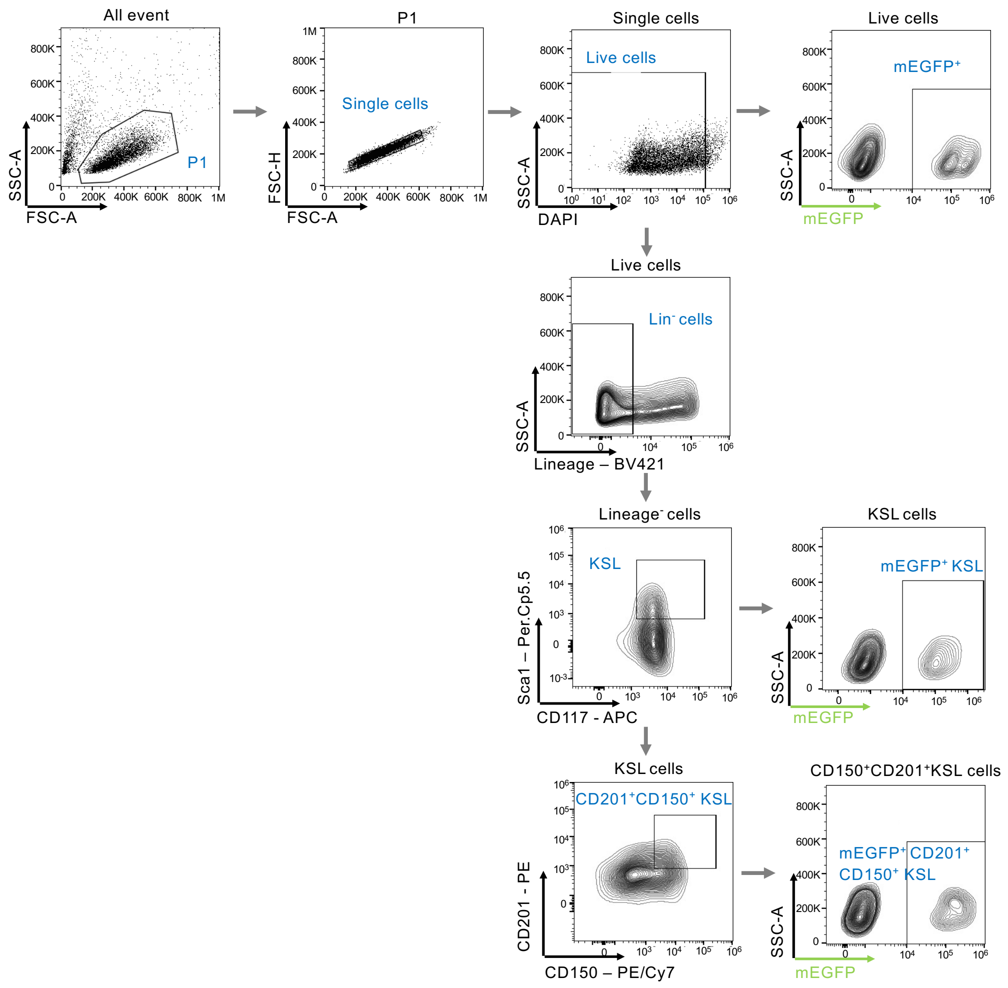


**Supplemental Figure 1.** Gating strategy for flow cytometry analysis to evaluate KI efficiency of *ex vivo* genome editing of mouse HSC.

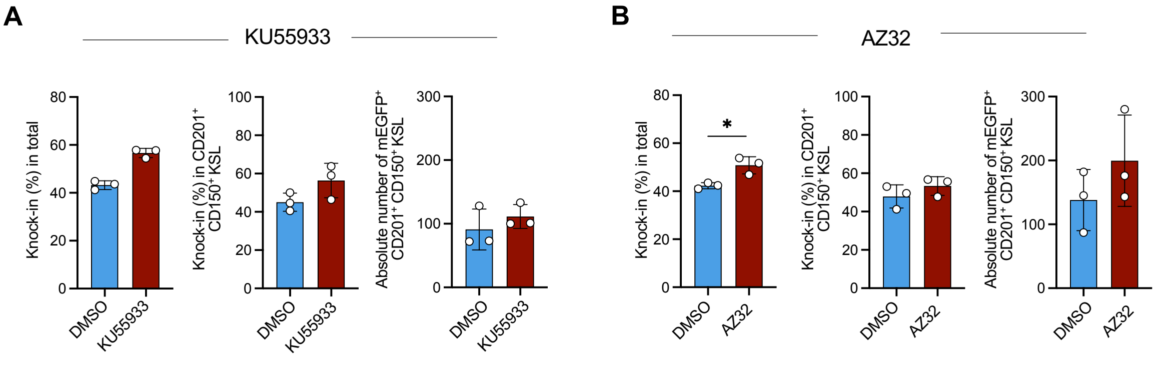


**Supplemental Figure 2.** Some of the ATM inhibitors tested for genome editing in mouse HSCs. (A) KU55933, (B) AZ32

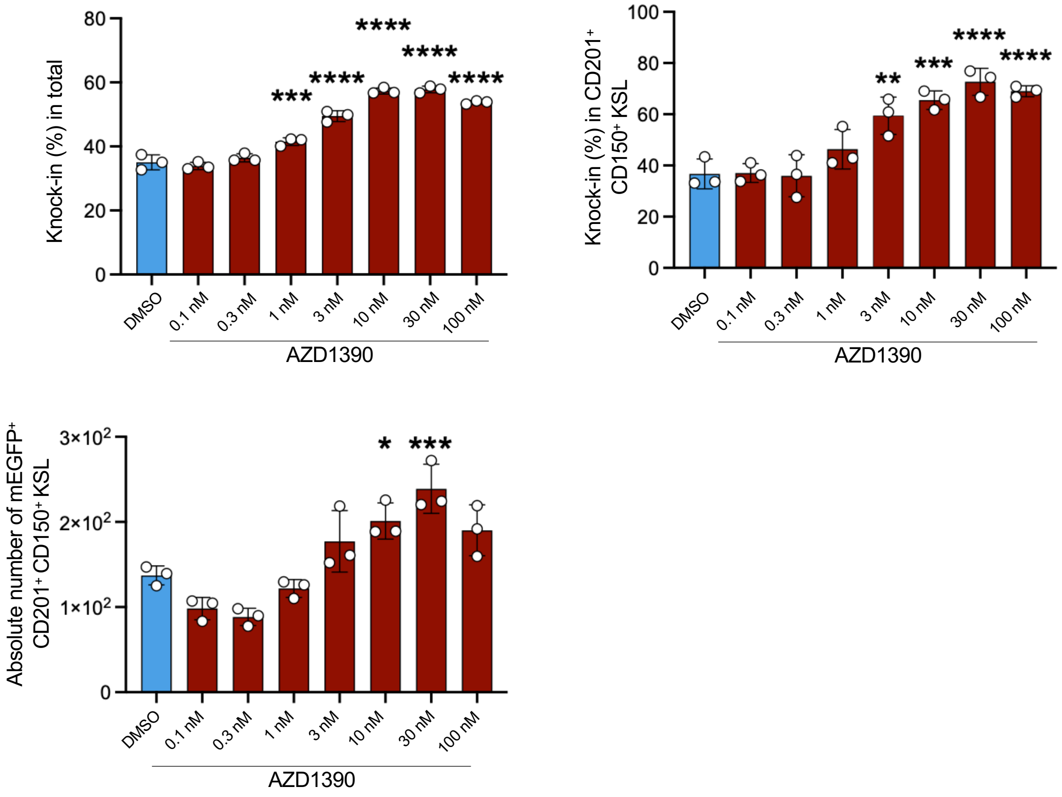


**Supplemental Figure 3**. AZD1390 dose-dependently increased the knock-in efficiencies in HSPC and HSC fraction.

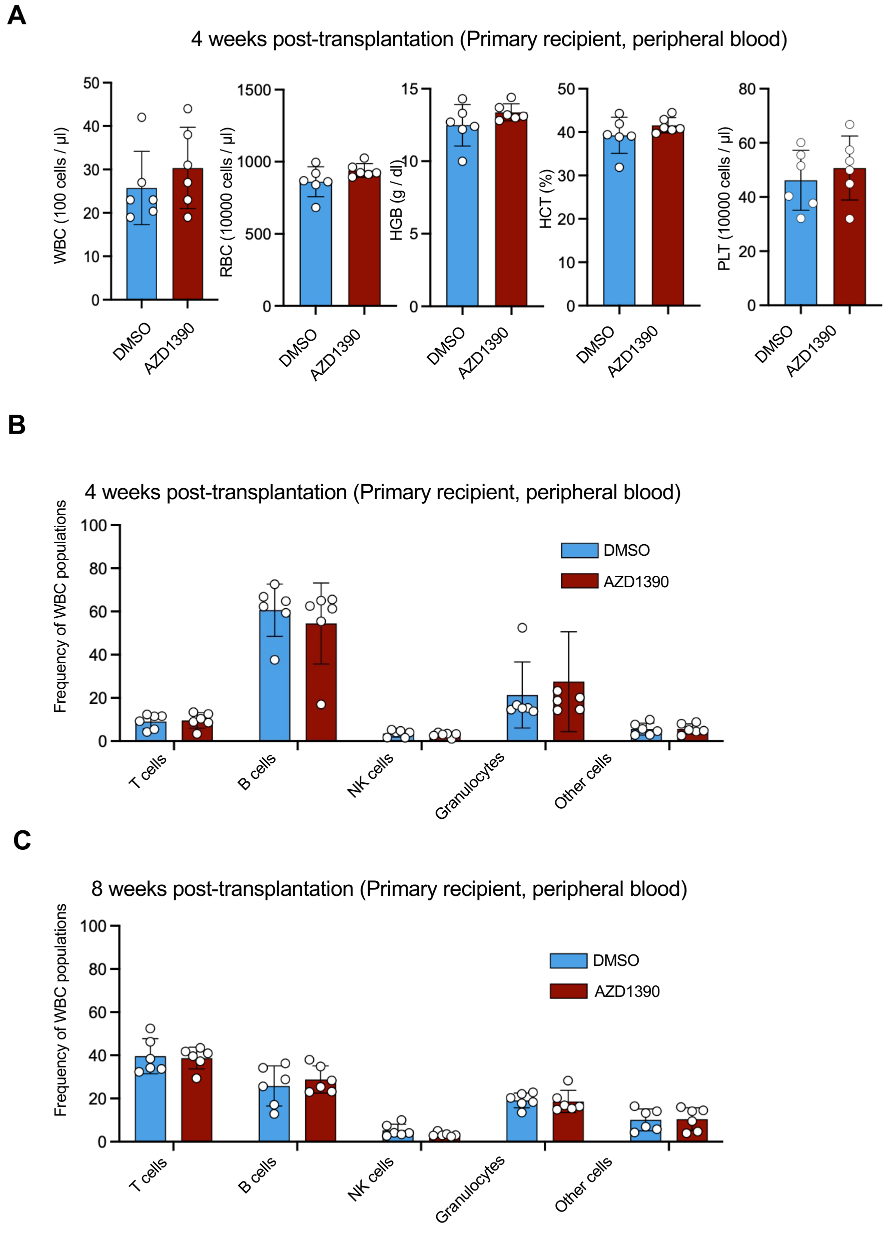


**Supplemental Figure 4**. CBC and PB WBC in recipient mice. (A) CBC results, (B-C) PB WBC

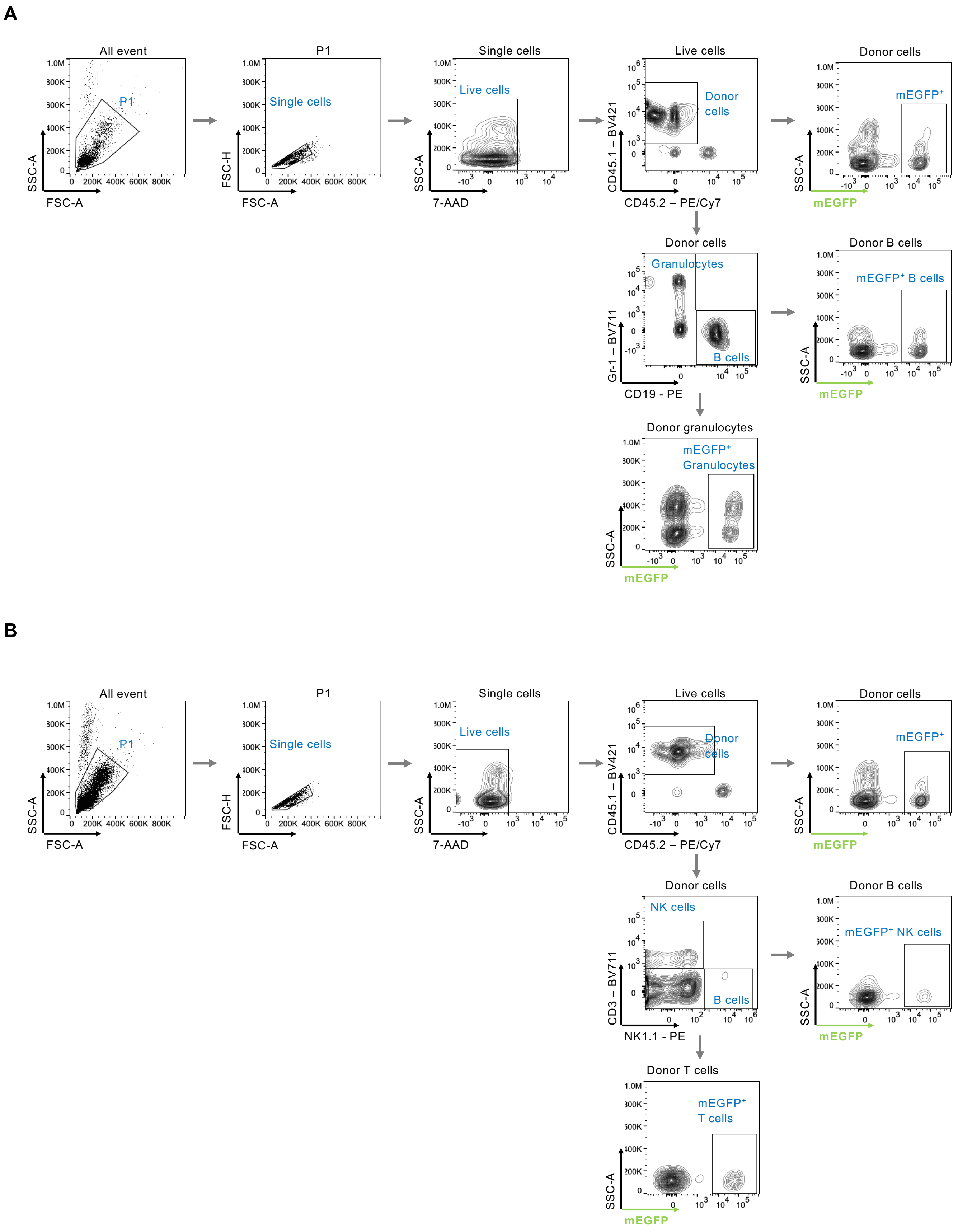


**Supplemental Figure 5.** Gating strategy for flow cytometry analysis to evaluate KI efficiency in PB WBCs. (A) Granulocytes and B cells, (B) T and NK cells.

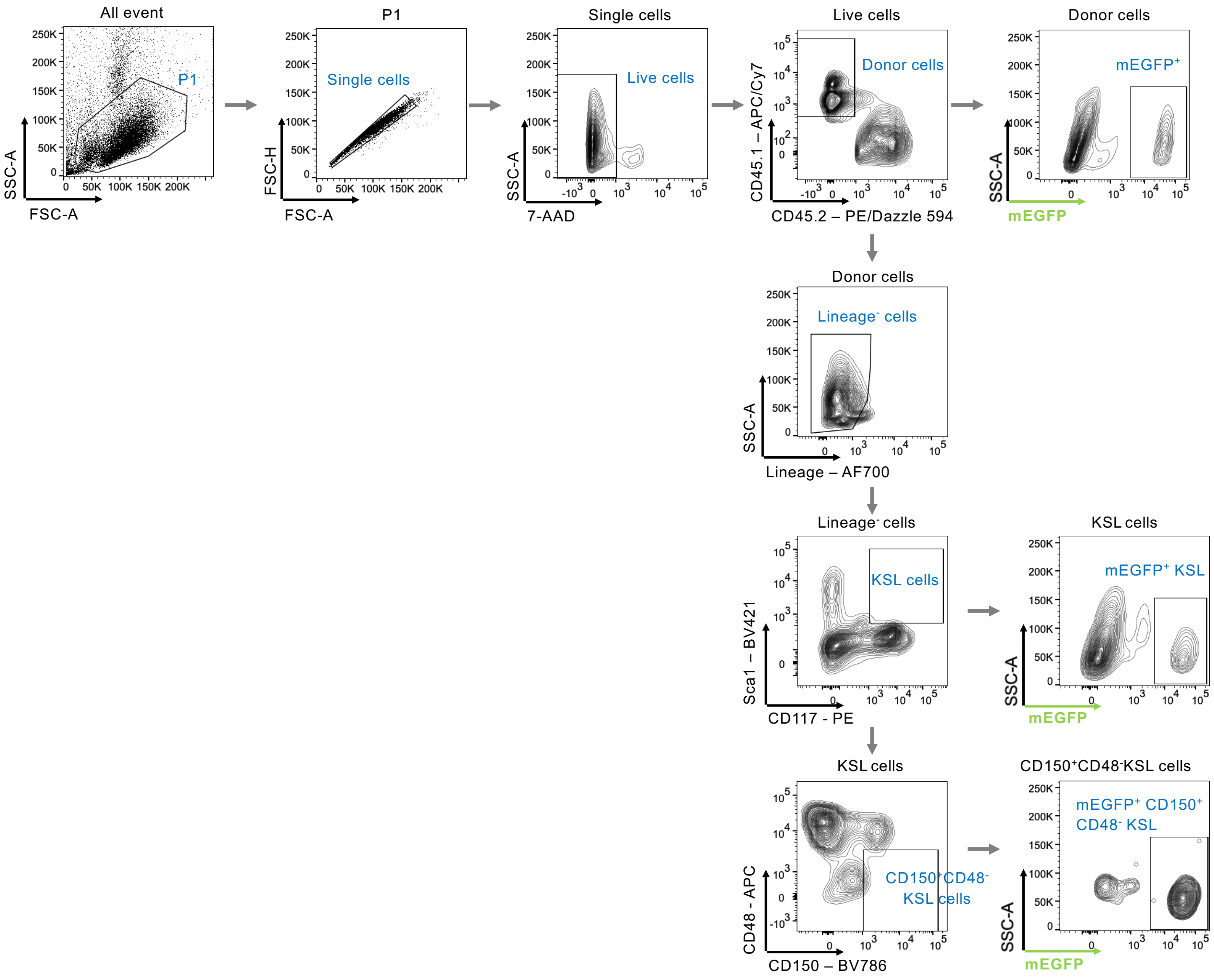


**Supplemental Figure 6.** Gating strategy for flow cytometry analysis to evaluate KI efficiency in BM cells.

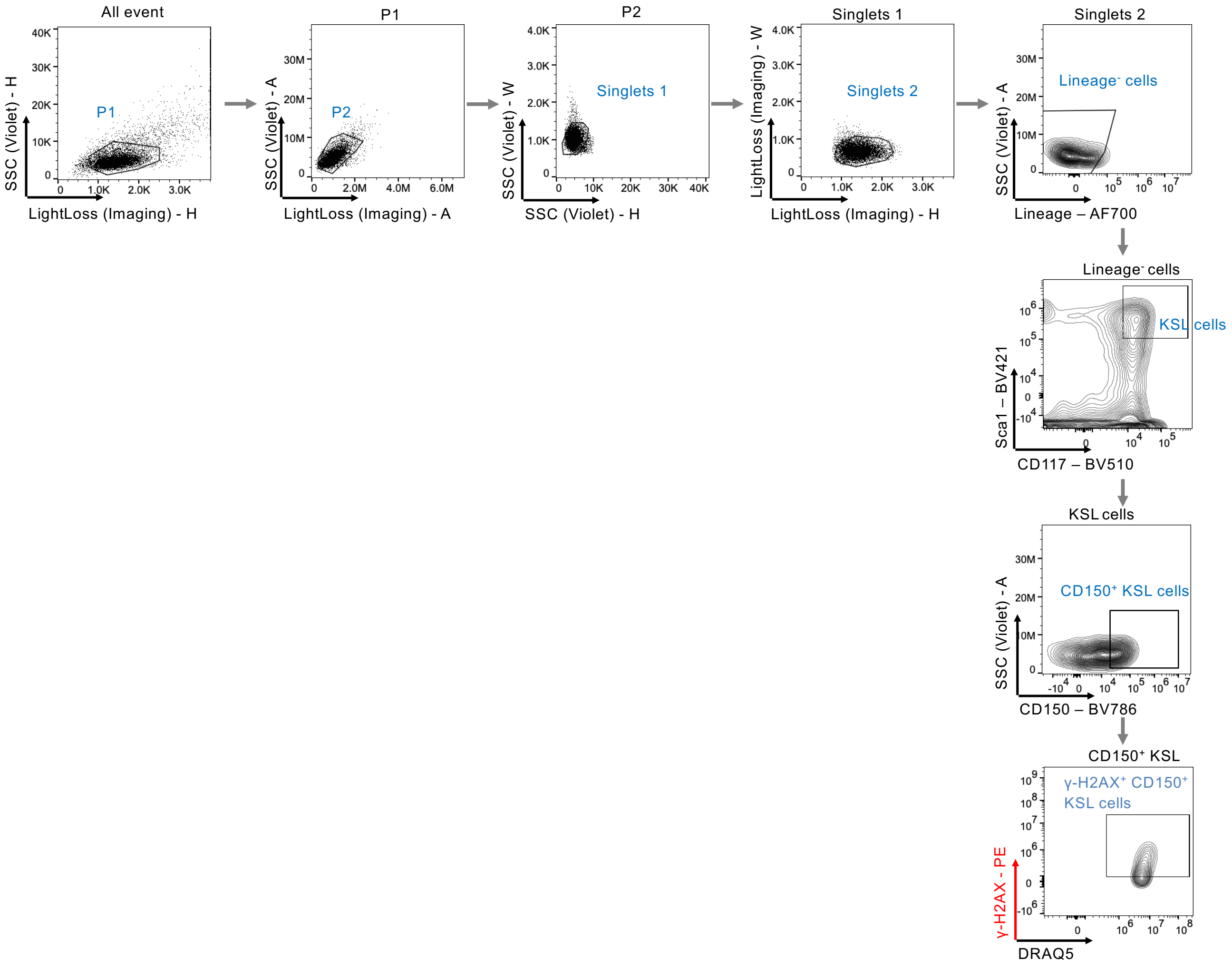


**Supplemental Figure 7.** Gating strategy for flow cytometry analysis to evaluate intracellular γ-H2AX level.

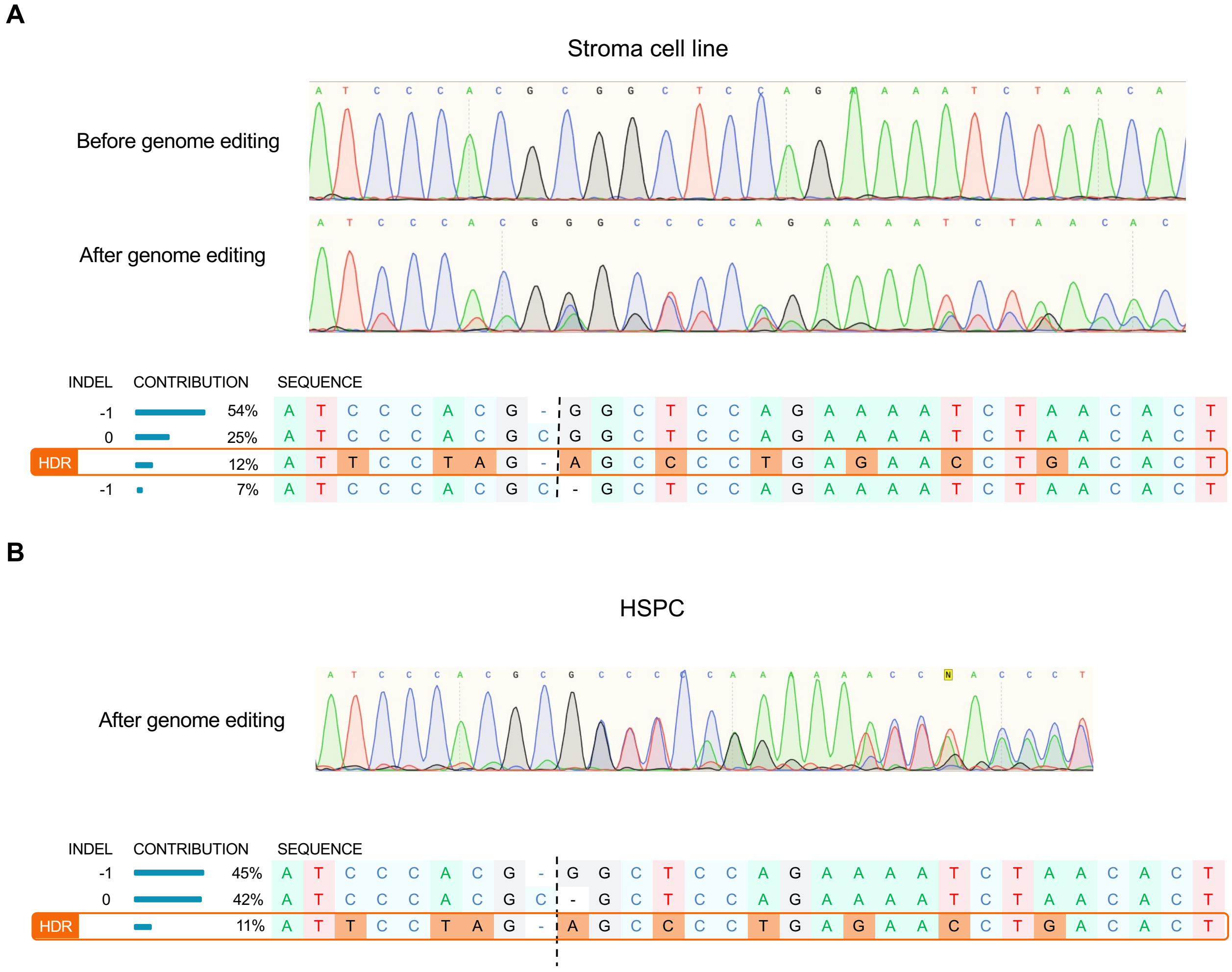


**Supplemental Figure 8.** Representative for sequencing results and analysis of HDR by sequencing trace file for *Il2rg* correction. (A) BM-derived stroma cells, (B) HSPCs.

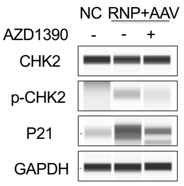


**Supplemental Figure 9.** ATM inhibition suppresses CHK2 and following p21 activation.
